# Tumor suppressor DAPK1 catalyzes Adhesion Assembly on Rigid but Anoikis on Soft Matrices

**DOI:** 10.1101/320739

**Authors:** Ruifang Qin, Haguy Wolfenson, Mayur Saxena, Michael P. Sheetz

## Abstract

Cancer cells will normally grow on soft surfaces, but if rigidity sensing modules are restored in cancer cells, they will undergo apoptosis on soft surfaces (*anoikis*) like most normal cells. DAPK1 is a major tumor suppressor that activates cell death, but it is unclear how DAPK1 could activate *anoikis* through rigidity sensing. Here we find that when rigidity sensing is decreased through inhibition of DAPK1 activity, cells are transformed for growth on soft matrices. Further, DAPK1 catalyzes matrix adhesion assembly and is part of adhesions on rigid surfaces. Additional factors include DAPK1 phosphorylation of tropomyosin2.1, talin1 head domain and tyrosine phosphorylation of DAPK1 by Src. On soft surfaces, DAPK1 rapidly dissociates from the adhesion complexes and activates apoptosis that requires PTPN12 activity and talin1 head. Thus, DAPK1 is important for adhesion assembly on rigid surfaces and the activation of *anoikis* on soft surfaces through its binding to rigidity-sensing modules.

## Introduction

In the early studies of cancer cells in vitro, it was evident that they could grow on widely different substrates, particularly on soft agar, which typically caused apoptosis of normal cells (a process called *anoikis*). This result indicated that either the cancer cells could not sense the surface properly, or that they were activated for growth under all conditions through receptor tyrosine kinases (RTK), Ras, or other mutations. Evidence for the former possibility came from recent studies showing that metastatic cancer cells (e.g., MDA-MB-231, HT1080 and SKOV3) undergo *anoikis* on soft agar after restoration of rigidity sensing modules. This was done through normalization of the expression levels of cytoskeletal proteins involved in rigidity sensing modules. Subsequent depletion of other proteins involved in the modules restored transformed growth^1^. However, the mechanism by which the rigidity sensing modules activated *anoikis* was unclear.

DAPK1 is a calcium⁄calmodulin-dependent protein kinase with cell death-inducing functions^2,3^. Not only is DAPK1 activity linked to *anoikis* but also it is a tumor suppressor that is depleted in many cancers^4–7^. Further, DAPK1 is linked to autophagy and apoptosis activated by TNF-alpha^8^. Classically, the activation of DAPK1 is believed to occur by calcium-calmodulin binding to DAPK1 to enable the dephosphorylation of the auto-inhibitory S308 site^9^. There is, however, additional evidence of inhibition through phosphorylation of vicinal tyrosines (Y490/Y491 in human DAPK1) and activation in immune cells by tyrosine phosphatase activity^10^. Still, no direct link between DAPK1 activity and rigidity sensing has been established. Some studies have linked DAPK1 to cytoskeletal proteins and motility, including the ability of DAPK1 to phosphorylate myosin light chain and tropomyosin2.1 (Tpm2.1)^11,12^ (formerly known as Tm1^13^). Notably, Tpm2.1 expression is sufficient to restore *anoikis* in several metastatic cancer lines^1,14^. Further, DAPK1 has a role in cell polarization and migration through an interaction with the talin1 head domain^15^. These observations raise the possibility that there may be a role for DAPK1 in rigidity sensing, as well as a role for rigidity sensing complexes in activating DAPK1 on soft surfaces.

Rigidity sensors in differentiated fibroblasts are relatively complicated protein modules that are capable of transducing matrix rigidity into biochemical signals to either activate growth on rigid matrices or apoptosis on soft matrices. The rigidity sensors are non-muscle sarcomeric-like units of about 2 μm in length, with anti-parallel actin filaments anchored to the matrix adhesions and a bipolar myosin IIA filament in the middle (reviewed in Wolfenson et al., Ann Rev Physiol In Press). To test the rigidity, these units contract the matrix to a constant distance of about 100 nm in total, and the force that is developed is sensed. Receptor tyrosine kinases, AXL and ROR2, regulate the displacement and the duration of the contractions, respectively^16^. EGFR and HER2 will activate contractions on rigid but not on soft matrices downstream of Src phosphorylation^17^. Surprisingly, normal rigidity sensing involves force-dependent cleavage of talin1 by calpain that catalyzes growth on rigid surfaces through the generation of free talin1 rod domains^18^. The velocity of the contraction depends upon Tpm2.1 binding to the actin filaments, and AXL phosphorylation of Tpm2.1 on Y214 is needed for adhesion formation^16^. Perhaps because this is a complicated sensory module, depletion of this module’s activity follows upon the depletion of any one of many cytoskeletal proteins, including Tpm2.1, myosin IIA, and α-actinin 4^19,20^. Loss of rigidity sensing activity, however, enables cell growth on soft surfaces, as is the case with many cancers. Upon restoration of rigidity sensing in cancer cells, many will die on soft surfaces, which begs the question of how does rigidity sensing activate apoptosis.

Here we studied the activation of *anoikis* through rigidity sensors, and found that DAPK1 activated *anoikis* and inhibition of DAPK1 was sufficient to enable growth on soft surfaces. Under control conditions, assembly of rigidity sensors and adhesion formation normally involved DAPK1 activity and particularly required the phosphorylation of Tpm2.1. On rigid substrates, DAPK1 associated with adhesions in a Tpm2.1-dependent process that involved Tpm2.1 phosphorylation by DAPK1. On soft substrates, DAPK1 rapidly dissociated from the adhesion complexes and activated apoptosis. Apoptosis was catalyzed by the constitutively phosphorylated mutant of Tpm2.1 and talin1 cleavage, but it was inhibited by Src and EGFR activity. DAPK1 assembly in adhesions depended on Y490/Y491 phosphorylation by Src or EGFR and tyrosine (Y) to phenylalanine (F) mutations catalyzed apoptosis. Inhibition of the tyrosine phosphatase, PTPN12, inhibited *anoikis* because it blocked DAPK1 activation. Thus, we suggest that DAPK1 is an important component in the assembly of rigidity sensors and adhesions through interactions with Tpm2.1 and talin1 head, leading to growth on rigid matrices and *anoikis* on soft matrices.

## Results

### Inhibition of DAPK1 contributes to cell growth on soft matrices

Because DAPK1 activity correlated with *anoikis*, autophagy, and TNF-alpha-induced apoptosis^8, 21, 22^, we wanted to test if it was responsible for apoptosis of cells on soft substrates. As shown in Fig. 1a, mouse embryonic fibroblasts (MEFs) on soft substrates (0.2 kPa) activated apoptosis with the characteristic changes in morphology, including membrane blebbing and cell rounding. Apoptosis of MEFs on soft substrates was blocked by inhibition of DAPK1 (Fig. 1a). Similar results were observed in MCF10A, normal human epithelial cells (Fig. 1a). Further quantitative analysis of cellular apoptosis based on Annexin V immunostaining matched cell apoptotic morphology results (Supplementary Fig. 1a). Thus, in both fibroblasts and epithelial cells, the cells were protected from apoptosis on soft matrices by an inhibitor of DAPK1.

**Fig. 1.**
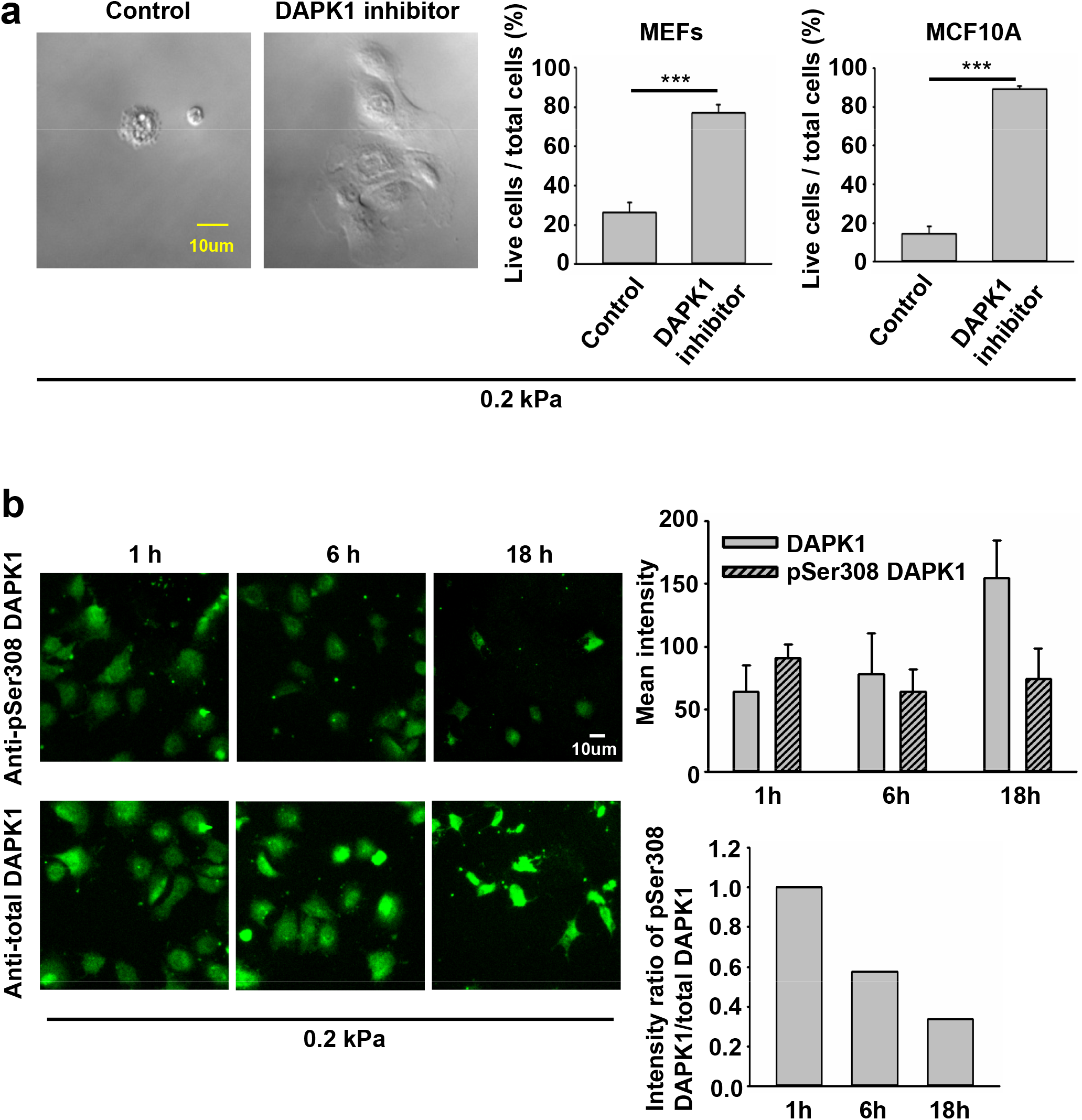
Inhibition of DAPK1 contributes to cell growth on soft matrices. (**a**) DIC images of MEFs and M10A cells with or without DAPK1 inhibitor after 2 days in culture on 0.2 kPa gels. Scale bar: 10 μm. Apoptotic cells were identified for appearance of apoptotic morphology including membrane blebbing and cell rounding. Live cells retained their normal spread morphology. Graphs represent means ± SEM of at least two independent experiments. ***P <0.001. (**b**) DAPK1-Ser308 phosphorylation was monitored by immunofluorescence in MEFs plated on 0.2 kPa gels. Immunofluorescence shows that DAPK1 was dephosphorylated at Ser-308 and thus activated on soft surfaces over time. Representative pictures are shown on the left panel for each condition. Scale bar: 10 μm. Graphs represent means ± SEM. Experiment was repeated twice.

The autophosphorylation of DAPK1 at S308 was found to be the major mechanism of DAPK1 inhibition in cells and that site was exposed for dephosphorylation upon binding of calcium/calmodulin^9^. To test if phospho-S308 decreased in *anoikis*, we followed the levels of pS308 by immunostaining with specific antibodies during the first 18 hours on soft surfaces. As predicted, the level of pS308 phosphorylation dropped on soft surfaces, indicating that DAPK1 was activated (Fig. 1b). Thus, there was activation of DAPK1 on soft surfaces with dephosphorylation of S308.

### DAPK1 regulates adhesion growth and force production

Inhibition of DAPK1 activity also slowed cell spreading and particularly adhesion formation (Fig. 2a-c). Further, the knockdown of DAPK1 by shRNA produced similar results (Supplementary Fig. 1b). Careful examination of the spreading process indicated that the early spreading phase was nearly normal without DAPK1 activity, but the later spreading phase that was associated with rigidity-sensing, contractile activity (Phase 2 of spreading^23^) was inhibited in correlation with decreased adhesion size. Thus, it seemed that DAPK1 activity was needed to form mature adhesions.

**Fig. 2.**
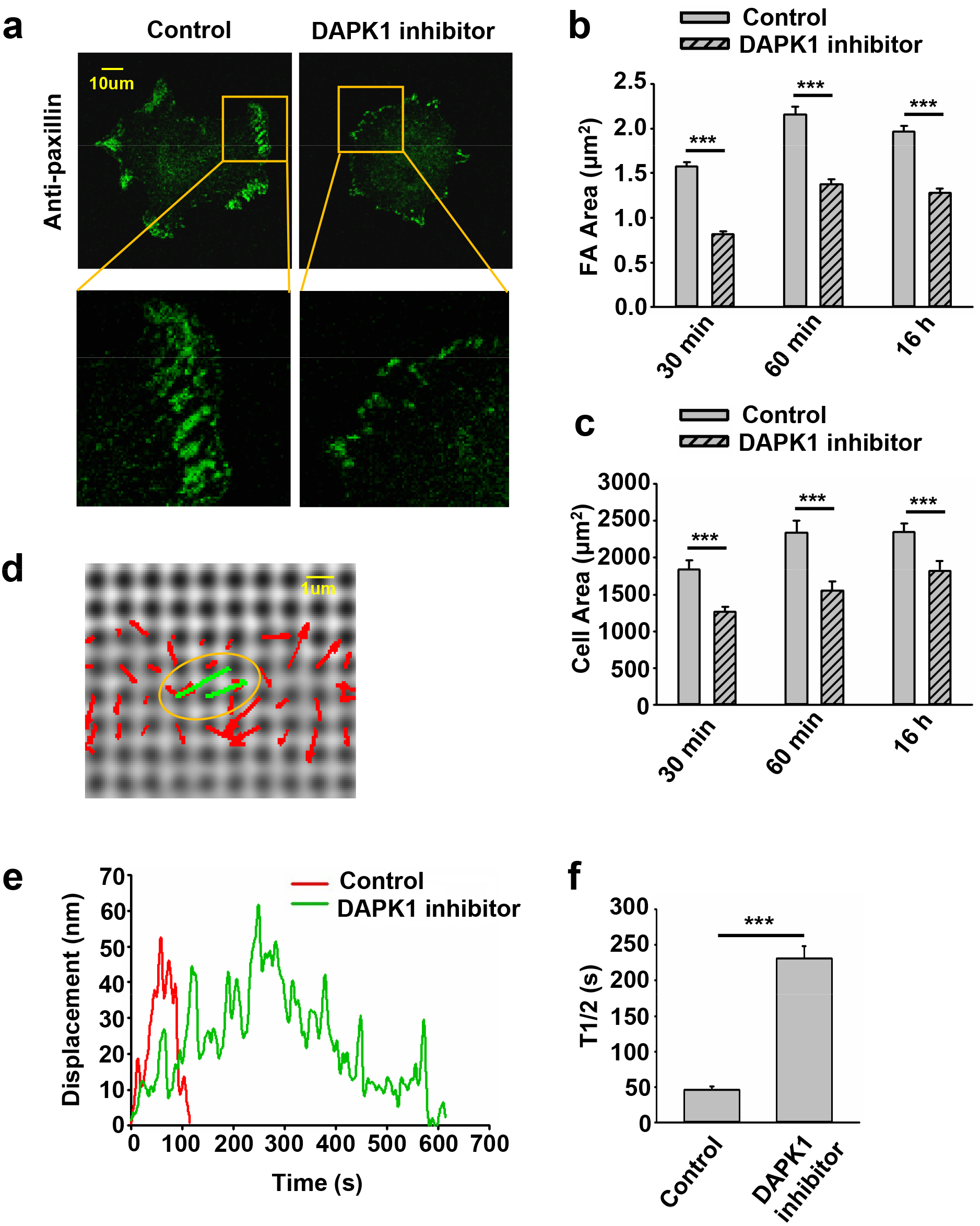
DAPK1 regulates adhesion growth and force production. (**a**) Micrographs showing the distribution of paxillin in MEFs treated with or without DAPK1 inhibitor. MEFs treated with or without DAPK1 inhibitor were fixed after spreading on fibronectin-coated glass dishes for 30 minutes, 60 minutes or 16 hours, followed by anti-paxillin/AlexaFluor 488 immunostaining. (**b**) The adhesion sizes were quantified (n>400 adhesions from >10 cells in each case, mean ± SEM). ***P <0.001. (**c**) Average area ± s.e.m of cells were also determined (n>10 cells in each case). ***P <0.001. Experiment was repeated three times. (**d**) Displacements (red arrows) of pillars that show the contractile pair (green arrows marked with yellow oval) at the periphery of a spreading MFF. Scale bar: 1 μm. (**e**) Displacements versus time of contractile pillars in MEFs treated with or without DAPK1 inhibitor. (**f**) Mean ± SEM of the T1/2 (time of contraction above half-maximal displacement) distributions of pillar displacements by MEFs before and after treatment with DAPK1 inhibitor. ***P < 0.001. N > 30 pillars in each case. Experiment was repeated twice.

We previously showed that local contractions to sense substrate rigidity correlated with adhesion formation and maturation^24^. To test if DAPK1 inhibition altered adhesion formation by affecting rigidity sensing contractions, cells were spread on fibronectin-coated 500 nm diameter PDMS pillars. When cells reached phase 2 of spreading, they contracted pairs of pillars toward each other to a peak distance of 60 nm each for about 30 seconds and then relaxed pillars at about the same velocity (2-3 nm/s), consistent with previous studies^19,24,25^. Inhibition of DAPK1 caused a dramatic decrease in the velocity of pulling (0.3-0.5 nm/s) with repeated release events (Fig. 2d-f), but did not alter the maximum pillar displacement (~60 nm) or contractile unit density (Supplementary Fig. 1c). Thus, the decrease in adhesion formation after DAPK1 inhibition correlated with a decrease in the rate of rigidity-sensing contractions.

### DAPK1 co-localizes with Tpm2.1 and targets to focal adhesion sites

Because the rigidity sensing contractions depended upon Tpm2.1^19^ and DAPK1, we then analyzed the distribution of DAPK1 and Tpm2.1 in early spreading of MCF10A epithelial cells. Both DAPK1 and Tpm2.1 colocalized at the spreading cell edges, indicating that they could be in a complex there (Fig. 3a). In Tpm2.1-transfected MDA-MB-231 cells (human breast cancer cell line) which normally lacked endogenous Tpm2.1^26^ but were rigidity-dependent for growth with Tpm2.1^1^, DAPK1 overlapped with Tpm2.1 at the cell edge. In contrast, further back from the edge, where Tpm2.1 was also present, decorating actin filaments, no overlap was observed (Fig. 3b). Thus, it seemed that Tpm2.1 and DAPK1 concentrated at active edge regions but not at all sites of Tpm2.1 concentration.

**Fig. 3.**
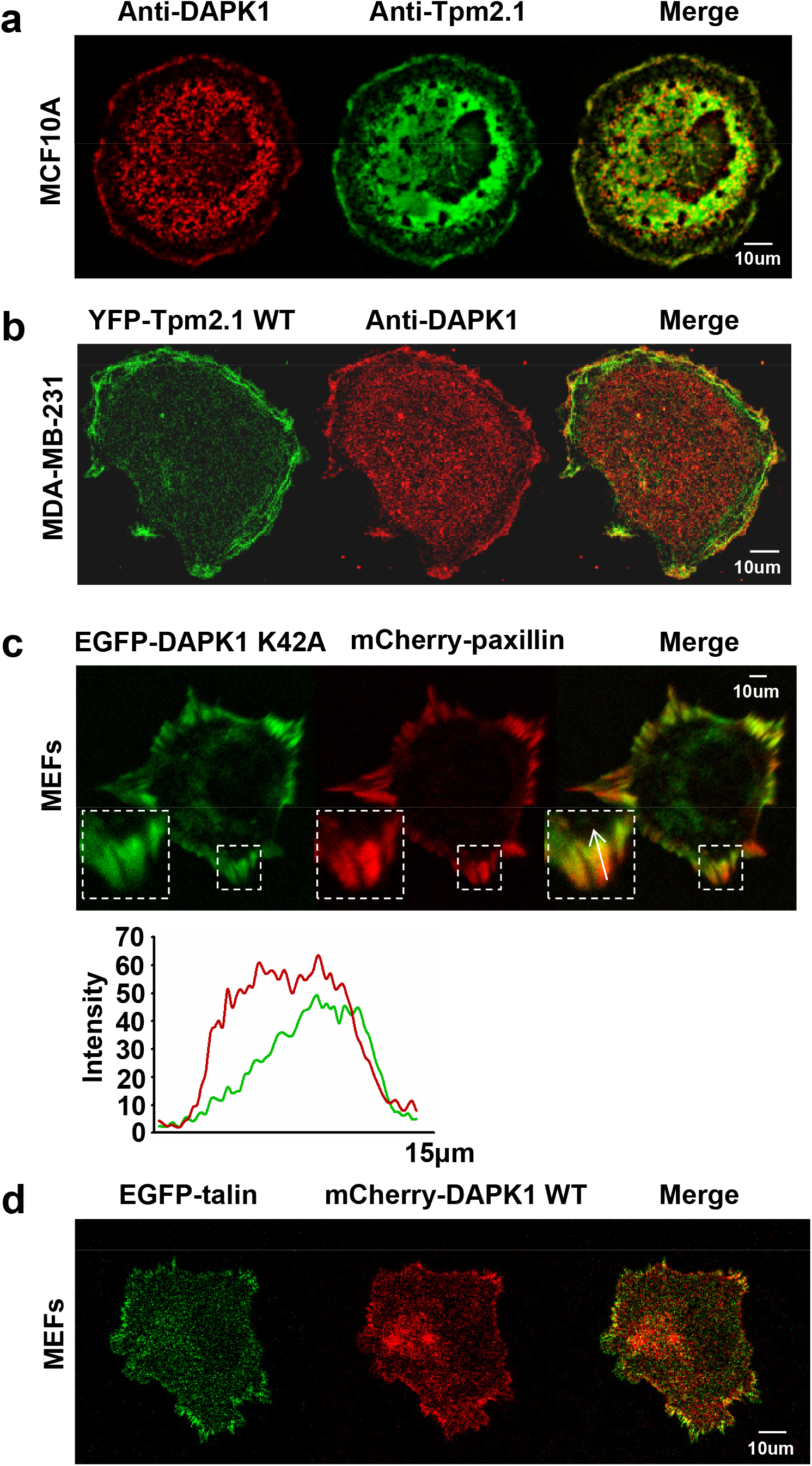
DAPK1 co-localizes with tropomyosin2.1 and targets to focal adhesion sites. (**a**) MCF10A cells were fixed after spreading on fibronectin-coated glass dishes for 30 minutes, followed by anti-tropomyosin2.1/AlexaFluor 488 and anti-DAPK1/AlexaFluor 555 immunostaining. Scale bar: 10 μm. (**b**) MDA-MB-231 cells were transfected with YFP-Tpm2.1 WT. Cells were further analyzed by anti-DAPK1/AlexaFluor555 immunostaining. Scale bar: 10 μm. (**c**) Total internal reflection fluorescence (TIRF) image of MEF co-transfected with EGFP-DAPK1 K42A and mCherry-paxillin after 30 minutes spreading on fibronectin-coated glass dish. Fluorescence intensity line profiles on the right are from the area of the representative adhesions covered by the white arrow in the merged images. Scale bar: 10 μm. (**d**) MEFs were co-transfected with EGFP-talin and mCherry-DAPK1 WT. Cells were subsequently allowed to spread for 30 minutes on fibronectin-coated glass dish. Scale bar: 10 μm.

To dynamically track DAPK1 in cells without causing cell blebbing and apoptosis due to its overexpression, we next used an EGFP-DAPK1 K42A construct (the inactive form of DAPK1)^27^ in MEFs. During early spreading of MEFs on fibronectin-coated glass, EGFP-DAPK1 K42A showed significant overlap with paxillin in focal adhesion sites, although the majority of DAPK1 was in the older portion of the adhesions (the edge closer to the nucleus) (Fig. 3c). We further confirmed in static images that DAPK1 WT (in the few cells that did not die) localized to focal adhesions like DAPK1 K42A (Fig. 3d; Supplementary Fig. 2a). Similar results were observed on fibronectin-coated pillars with either the DAPK1 construct or anti-DAPK1 antibody (Supplementary Fig. 2b, c). These results all indicated that DAPK1 was part of adhesions on fibronectin-coated surfaces in cells that were rigidity-dependent for growth.

### The phosphorylation of Tpm2.1 by DAPK1 is important for adhesion formation but increases sensitivity to *anoikis*

Since DAPK1 was found to phosphorylate Tpm2.1 on S283^11^, the inhibition of DAPK1 activity may have altered cell adhesion formation through the inhibition of Tpm2.1 phosphorylation. To test this possibility, we transfected MDA-MB-231 cells with mutant forms of Tpm2.1 which were blocked for phosphorylation, Tpm2.1 S283A, or mimicked the phosphorylated form, Tpm2.1 S283E. Adhesion size depended upon Tpm2.1 transfection and with the non-phosphorylatable mutant, small adhesions formed that were similar in size to non-transfected MDA-MB-231 cells. In contrast, with the Tpm2.1 S283E mutant, larger adhesions formed and cells spread to larger areas similar to Tpm2.1 WT (Fig. 4a, b). In the Tpm2.1 S283A-transfected cells, there was increased resistance to apoptosis on soft surfaces, whereas the Tpm2.1 S283E cells had greater sensitivity to apoptosis on soft surfaces than Tpm2.1 WT transfected ones (Fig. 4c). Thus, the phosphorylation of Tpm2.1 at the DAPK1 phosphorylation site was important for adhesion formation in MDA-MB-231 cells but facilitated *anoikis* on soft substrates.

**Fig. 4.**
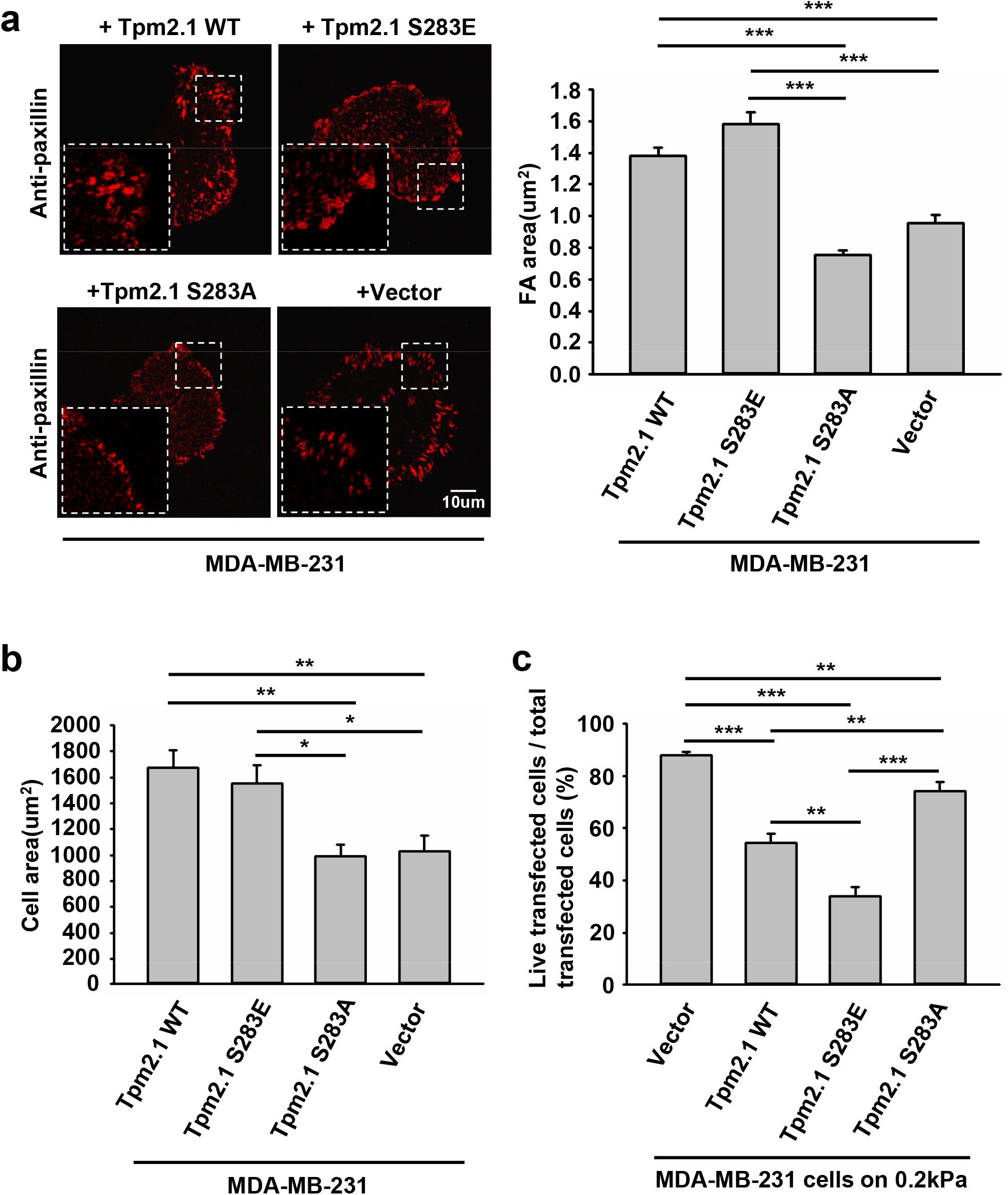
The phosphorylation of Tpm2.1 by DAPK1 is important for adhesion formation but facilitates *anoikis* on soft substrates. (**a**) Left: micrographs showing the distribution of paxillin in MDA-MB-231 cells transfected with YFP-Tpm2.1 WT, YFP-Tpm2.1 S283E (phospho-mimic mutant), YFP-Tpm2.1 S283A (non-phosphorylated mutant) or YFP-Vector respectively after 30 minutes spreading on fibronectin-coated glass dishes, followed by anti-paxillin/AlexaFluor 555 immunostaining. Scale bar: 10 μm. Right: the adhesion sizes were quantified (n>400 adhesions from >10 cells in each case, mean ± SEM), ***P <0.001. (**b**) Average area ± s.e.m of transfected cells in each group were determined (n>10 cells in each case). *P <0.05, **P <0.01. (**c**) MDA-MB-231 cells were transfected with YFP-Tpm2.1 WT, YFP-Tpm2.1 S283E, YFP-Tpm2.1 S283A or YFP-Vector respectively for one day and replated on fibronectin-coated 0.2 kPa gels for one day. The percentage of transfected cells that were still alive was quantified. The mean ± SEM of at least two independent experiments is described. For each experiment, 100∼150 cells were analyzed for each transfection point. **P <0.01, ***P <0.001.

### Talin1 head aids in the localization of DAPK1 to the adhesions and facilitates apoptosis

Previous studies reported that DAPK1 interacted with talin1 through the head domain^15^. In a separate study, we also showed that the assembly of adhesions depended upon talin1 cleavage by calpain and cell growth depended upon the presence of the talin1 rod in the absence of the head^18^. To test if talin1 cleavage was needed for DAPK1 assembly into adhesions, we transfected non-cleavable (NC) talin1, WT talin1 and the rod and head fragments separately in talin1-/- fibroblasts. With WT talin1 and the talin1 head, we found that DAPK1 assembled into focal adhesions; however, neither the talin1 rod nor NC talin1 supported adhesion assembly with DAPK1 (Fig. 5a). Thus, although the rod was needed for growth, the head fragment was needed for DAPK1 assembly in adhesions, which supported previous findings of an interaction of DAPK1 with the talin head^15^. In DAPK1 WT transfected talin1-/- cells with talin1 constructs or MEFs treated with a calpain inhibitor (to prevent talin cleavage), talin1 head and talin1 cleavage caused an increase in DAPK1 apoptotic function (Fig. 5b, c). When talin1-/- cells were cultured on soft surfaces, they were protected against apoptosis whereas with the WT talin1 they apoptosed on soft surfaces. Rescue of talin1-/- cells with NC-talin1 did not restore apoptosis on soft surfaces but expression of the talin1 head domain caused greater apoptosis on soft surfaces than did WT talin1 (Supplementary Fig. 3). Thus, the cleavage of talin1 by calpain produced the talin1 head domain fragment that was needed for DAPK1 to assemble in focal adhesions and to catalyze apoptosis on soft surfaces.

**Fig. 5.**
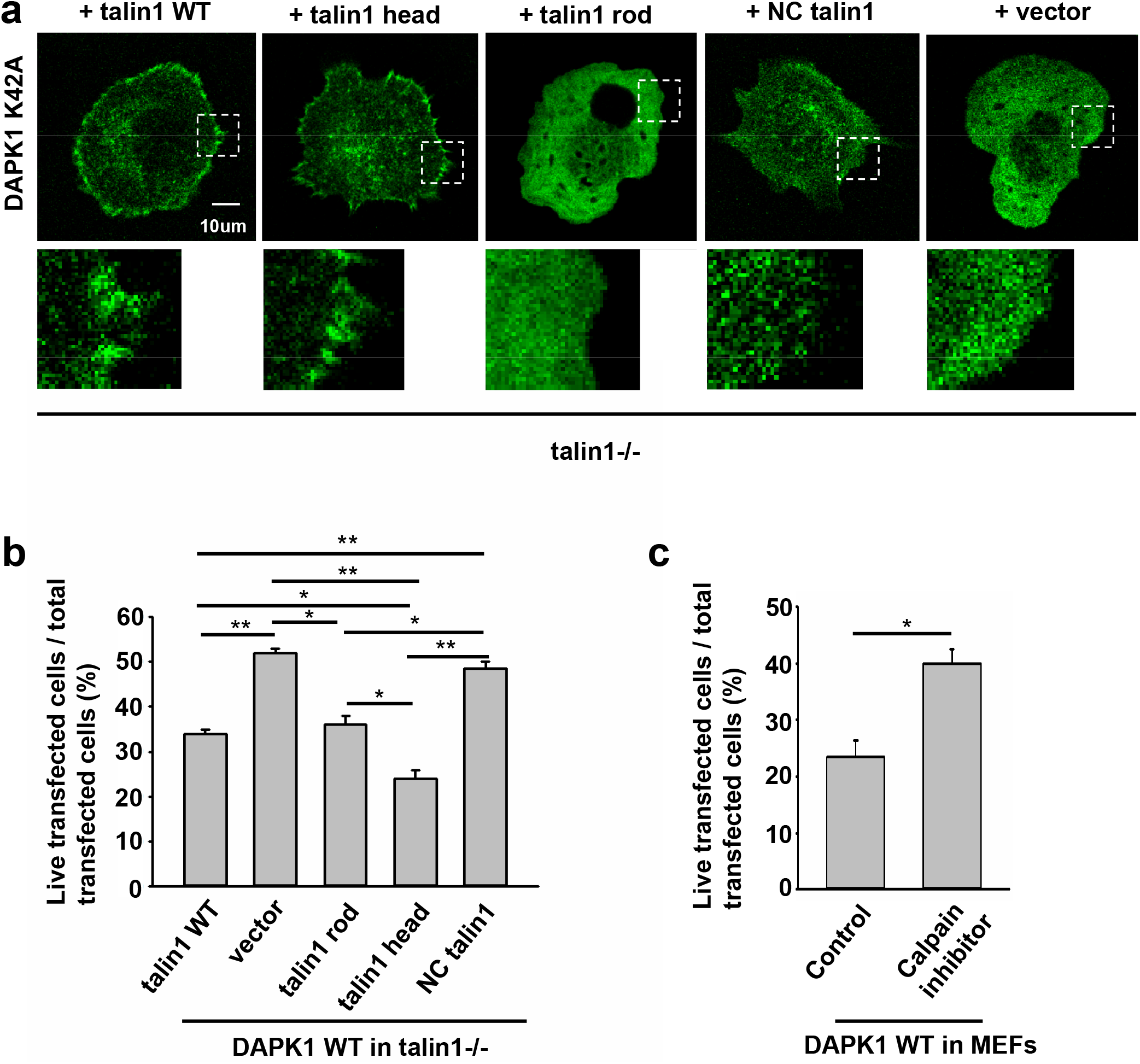
Talin1 head aids in the localization of DAPK1 to the adhesions and facilitates apoptosis. (**a**) Talin1-/- cells were co-transfected with DAPK1 and talin1 constructs (talin1 WT, talin1 head, talin1 rod, non-cleavable talin1) or vector. Cells were spread on fibronectin (30 minutes). Scale bar: 10 μm. (**b**) Talin1-/- cells were co-transfected with DAPK1 WT and talin1 constructs (talin1 WT, talin1 head, talin1 rod, NC talin1) or vector for one day. The percentage of co-transfected cells that were still alive was quantified (mean ± SEM of ≥2 experiments with 100∼150 cells; *P <0.05, **P <0.01) (**c**) MEFs transfected with EGFP-DAPK1 WT treated with DMSO or calpain inhibitor ALLN (100 μM). The percentage of transfected cells that were still alive was quantified. (mean ± SEM of ≥2 experiments with 100∼150 cells; *P <0.05).

### Dynamics of DAPK1 during cell spreading on dual stiffness pillars

To confirm that DAPK1 recruitment to adhesions was dependent upon local rigidity, we prepared pillar substrates that had sub-cellular regions with a 20-fold difference in pillar rigidity by exposing PDMS pillars to UV light in localized regions^28^. When MEF cells transfected with EGFP-DAPK1 K42A and mCherry-paxillin were spread on those dual stiffness pillars for 30 minutes, both DAPK1 and paxillin staining was lower on the soft than the rigid pillars (Fig. 6a). In time-lapse studies of DAPK1 recruitment to pillars, EGFP-DAPK1 K42A accumulated equally rapidly on soft and stiff pillars at early times, but was then released from the soft pillars after 1-2 minutes whereas it remained associated with the rigid pillars (Fig. 6b, c). We then followed single pillars during the contraction and relaxation to correlate the recruitment level of DAPK1 with contractile unit force. After the peak of contraction force, DAPK1 intensity on the soft pillars decreased rapidly (Fig. 6d). Thus, the recruitment of DAPK1 to adhesions was rapid but there was also a rapid release from soft pillars. To further determine if DAPK1 recruitment to or release from soft pillars was altered, we measured photobleaching recovery of EGFP-DAPK1 at soft and rigid pillars. The recovery rates were the same on both, indicating that binding was unaltered but release was accelerated on soft surface (Supplementary Fig. 4a, b). Thus, it seems that soft matrices cause the rapid release of DAPK1 from adhesion sites but recruitment continues.

**Fig. 6.**
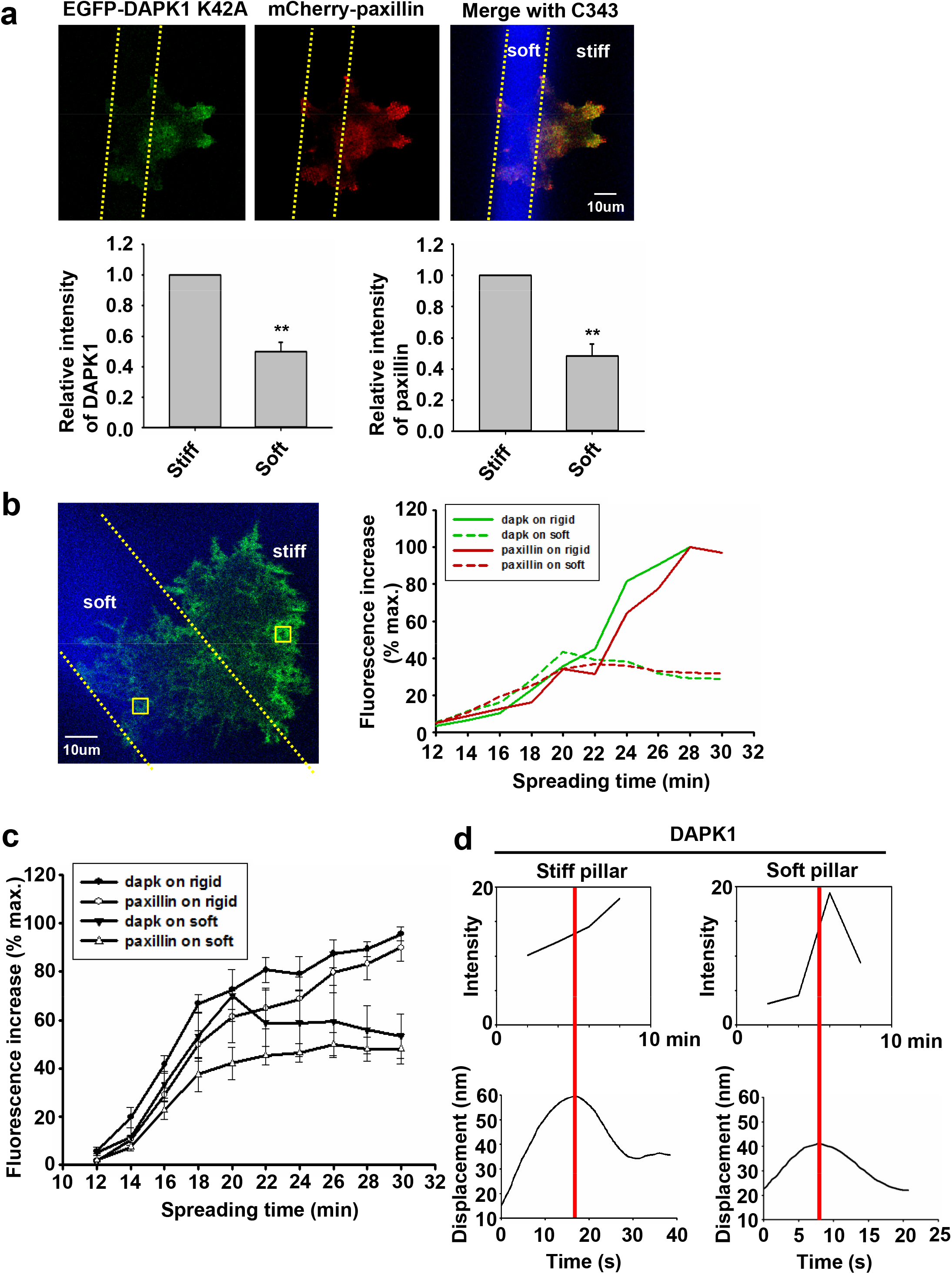
Dynamics of DAPK1 during cell spreading on dual stiffness pillars. (**a**) MEFs were co-transfected with EGFP-DAPK1 K42A and mCherry-paxillin. Cells were spread (30 minutes) on dual stiffness pillars (9kPa (Coumarin 343) and 170kPa), Scale bar: 10 μm. The leading edge protein relative intensity was normalized by dividing the average intensity value within 3 micrometers of the cell edge in stiff region or soft region by the average intensity value of the cell edge in stiff region of each cell (stiff /stiff or soft /stiff). The relative intensity of DAPK1 or paxillin is presented in the lower panels. **P <0.01. N=at least 5 different cells in each group. Graphs represent mean ± SEM. (**b**) Left: confocal image showing a cell transfected with EGFP-DAPK1 K42A and mCherry-paxillin on dual stiffness pillars. Scale bar: 10 μm. Right: one adhesion site of the left cell corresponding to a single pillar in a stiff or soft region was tracked during cell spreading. The tracked pillar is marked by a yellow box. The relative fluorescence of both molecules was plotted. (**c**) Quantification of relative increase in DAPK1 and paxillin fluorescence over time on dual stiffness pillars within 3 micrometers of outward curving cell edges. Four cells were measured on 2 different days. Graphs represent means ± SEM. (**d**) The time course of recruitment of DAPK1 was measured during pillar contraction and relaxation. Displacements versus time of contractile pillars are shown in the lower panel. The peak of the pillar contraction force is marked by a red line.

### DAPK1 apoptotic function is controlled by Src phosphorylation and PTPN12

In previous studies, tyrosine kinase/phosphatase activities were implicated in the regulation of DAPK1^10^. To test if the vicinal tyrosines in DAPK1 (Y490/Y491) were involved, we overexpressed EGFP-DAPK1 mutants Y490D/Y491D (DYD, phospho-mimic mutant), Y490F/Y491F (DYF, non-phosphorylated mutant) or Y490F/Y491F-K42A (DYF-K42A, non-phosphorylated inactive mutant) in MEFs. Most of the Y-D mutation transfected cells (>80%) had concentrations of DAPK1 in adhesion sites, whereas there was much less DAPK1 assembly in adhesion sites with the DYF (∼30%) or DYF-K42A (∼20%) transfected cells caused (Fig. 7a). In an apoptosis assay, the Y-F mutation enabled normal apoptosis whereas the Y-D mutation inhibited apoptosis as did the Y-F double mutation of the inactive DAPK1-K42A (Fig. 7b). To test for the possible involvement of the Src family tyrosine kinases (SFK), we added the general SFK inhibitor PP2 and found that there was less DAPK1 assembled into adhesions (Supplementary Fig. 5a, b), as well as a dramatic decrease in adhesion size as had been reported before because of the inhibition of rigidity sensing contractions^17^. We next tested for apoptosis in the presence of PP2 and found a dramatic increase in DAPK1-induced cell death (Supplementary Fig. 5c). Since the SFKs also activated EGFR in early spreading^17^, we tested for the effect of EGFR inhibition on apoptosis and found enhanced cell death after EGFR inhibition (Supplementary Fig. 5c). Thus, tyrosine phosphorylation of DAPK1 on the vicinal tyrosines enabled assembly of DAPK1 into adhesions and inhibited apoptosis whereas the unphosphorylated DAPK1 couldn’t assemble in adhesions but supported apoptosis.

**Fig. 7.**
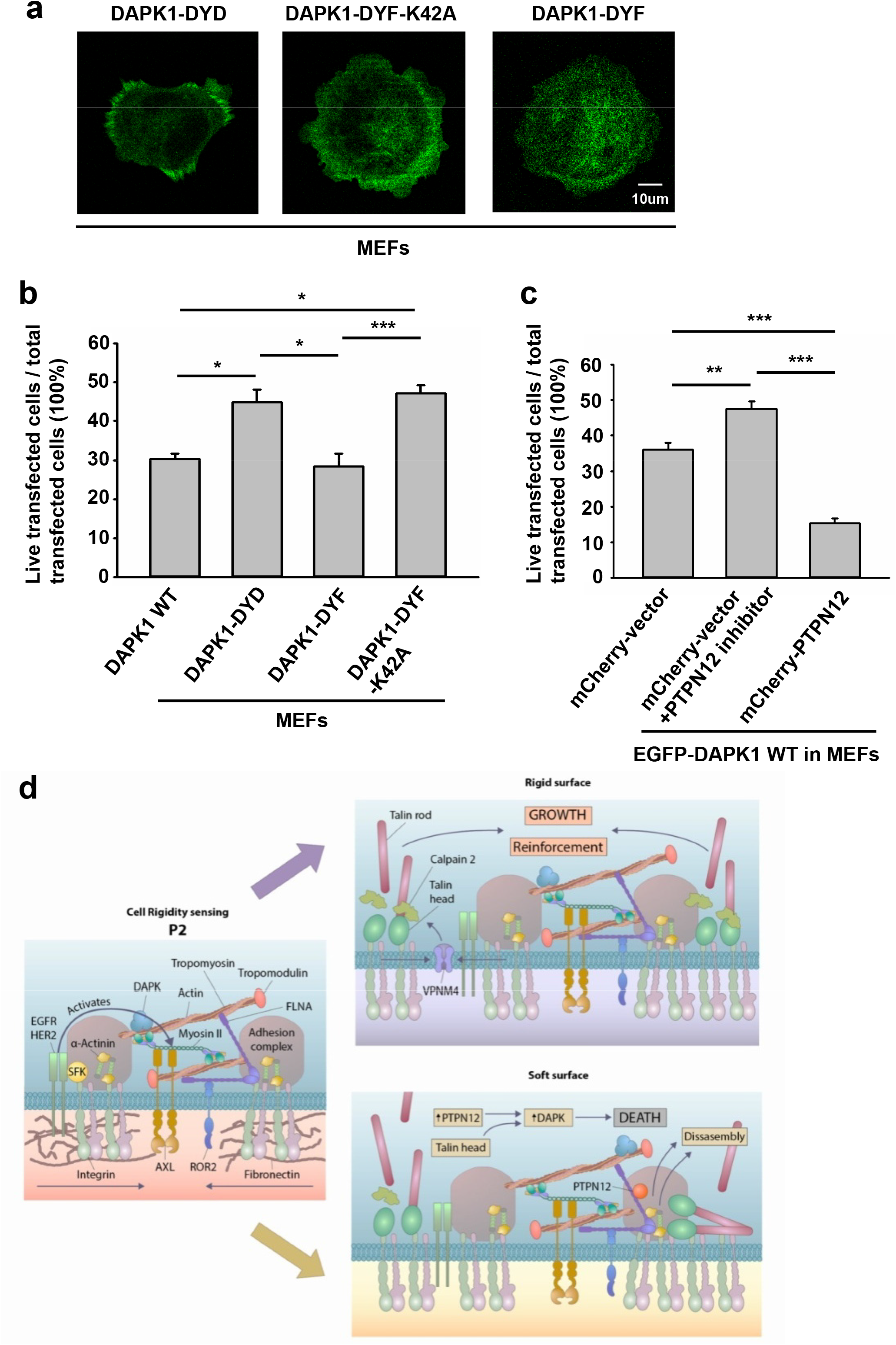
DAPK1 apoptotic activity is controlled by Src phosphorylation and PTPN12. (**a**) Micrographs showing the distribution of DAPK1 in MEFs transfected with EGFP-DAPK1-DYD (phospho-mimic mutant), EGFP-DAPK1-DYF (non-phosphorylated mutant) or EGFP-DAPK1-DYF-K42A (non-phosphorylated inactive mutant) respectively after 30 minutes spreading on fibronectin-coated glass dishes. Scale bar: 10 μm. (**b**) MEFs were transfected with EGFP-DAPK1 WT, EGFP-DAPK1-DYD, EGFP-DAPK1-DYF or EGFP-DAPK1-DYF-K42A respectively for one day. The percentage of transfected cells that were still alive was quantified. The mean ± SEM of at least two independent experiments is described. For each experiment, 100∼150 cells were analyzed for each transfection point. *P <0.05, ***P <0.001. (**c**) MEFs were transfected with EGFP-DAPK1 WT either co-transfected with mCherry-PTPN12 or treated with PTPN12 inhibitor. The percentage of co-transfected cells that were still alive was quantified. The mean ± SEM of at least two independent experiments is described. For each experiment, 100∼150 cells were analyzed for each transfection point. **P <0.01, ***P <0.001. (**d**) Schematic of a working model of DAPK1 shuttling depending on substrate rigidity. The presence of a rigid substrate keeps DAPK1 at the FA through mechanical tension. In contrast, a soft substrate induces DAPK1 disassembly from FA and activates DAPK1 apoptotic function.

There was reason to believe that DAPK1 assembly into the adhesions was a normal part of the cell-matrix interaction and that a phosphatase was needed to disassemble adhesions on soft surfaces and to activate DAPK1. A major tyrosine phosphatase that was implicated in cancer and motility was PTPN12^29,30^. To test for its possible involvement, we added an inhibitor of PTPN12 and found that it blocked DAPK1 apoptotic function in MEFs. Further, we observed an enhancement of apoptosis if we overexpressed PTPN12 (Fig. 7c). Thus, it seemed that tyrosine phosphorylation inhibited DAPK1 and apoptosis whereas PTPN12 activity was needed for activation of apoptosis on soft surfaces. These findings were summarized in a model diagram (Fig. 7d).

## Discussion

These studies show that DAPK1 has a major role both in the formation of matrix adhesions on rigid surfaces at early times and in the apoptosis of cells on soft surfaces. Both processes involve DAPK1 assembly in adhesions through its phosphorylation of Tpm2.1 at S283, as well as the proteolysis of talin1 to produce the head fragment. DAPK1 activity is needed for normal assembly of focal adhesions and proper rigidity sensing contractions. Further, the phosphorylation of DAPK1 on the vicinal tyrosine residues (Y490/491) by Src or EGFR is critical for its inactivation at adhesions. On sub-micrometer pillars with sub-cellular areas of different stiffness, the binding of DAPK1 to both soft and rigid pillars is initially similar, but once a maximal force is reached, rapid release is only found in the soft regions. DAPK1 activation, release, and adhesion disassembly all appear to correlate with the dephosphorylation of DAPK1 by PTPN12. Thus, the rigidity dependent activation of apoptosis involves DAPK1 activity in the formation of adhesions through Tpm2.1 and rigidity-dependent dephosphorylation of DAPK1.

An earlier study found that DAPK1 had a role in cell migration and polarization through an interaction with integrin and the talin1 head domain^15^. This is consistent with its function in the formation of contractile adhesions and rigidity sensing; however, the longer-term reactions to rigid surfaces involve activation of motility responses beyond early rigidity sensing. Actin-dependent membrane extension correlates with rigidity sensing and fully spread cells have few rigidity-sensing contractions because they have few extensions^17^. The cycle of mechano-testing and response can result in migration if there is polarization with a single leading edge. In the case of transformed cancer cells, there is a major change in cell state since they lack rigidity sensing contractions and Tpm2.1 in adhesions. It seems that normal rigidity sensing involves the recruitment of tyrosine phosphorylated DAPK1 along with Tpm2.1 to the contractile units. If the surface is rigid, then endogenous DAPK1 is stabilized in the adhesions along with Tpm2.1, talin1 head, and other focal adhesion proteins. If the surface is soft, then a phosphatase, such as PTPN12, dephosphorylates DAPK1 and causes its activation as well as the disassembly of the adhesions. In studies of immune cells, the phosphatase LAR dephosphorylates DAPK1 but that is not related to cell mechanics^10^. The role of PTPN12 in activating *anoikis* is consistent with its role as a tumor suppressor^30^ and it also has major effects on adhesion lifetime^31^.

The role of the tyrosine kinases in inactivating DAPK1 is consistent with previous studies that show high levels of phosphorylation of the vicinal tyrosine residues in cancer as well as fibroblasts^10^. In these studies, DAPK1 is inactivated upon phosphorylation by Src either directly or downstream through EGFR activation. In earlier studies, we found that Src activated EGFR or HER2 in cell spreading to catalyze rigidity sensing^17^. Here we find that inhibitors of either will cause increased apoptosis, indicating that DAPK1 normally is phosphorylated in the adhesions. Further, the inhibition of PTPN12 will block DAPK1 activation. The level of pS308 decreases on soft surfaces over time and the level of DAPK1 expression appears to increase as observed in other studies of DAPK1 activation but that may be through continued expression without assembly into adhesions on the soft surfaces. There are few rigidity sensing contractions on soft surfaces and few chances for assembly of DAPK1 into adhesion complexes with tyrosine phosphorylation^17^.

It is logical that the activation of apoptosis on soft surfaces would involve engagement of the death-associated protein kinase with the rigidity sensing process at integrin adhesions. Rigidity sensing involves the assembly of sarcomere-like contractile units at integrin adhesion sites and the subsequent pulling of those adhesions to produce a displacement of 100 nm. If the force developed is less than about 25 pN, then the matrix is considered soft and the adhesions disassemble, releasing activated DAPK1. Based upon these findings we hypothesize that Tpm2.1 and talin1 head domain form a complex with tyrosine phosphorylated DAPK1 that dissociates on soft surfaces due to the tyrosine phosphatase, PTPN12 (Fig. 7d).

The complex nature of the rigidity sensing module indicates that it integrates mechanosensing with several output signals depending upon the matrix rigidity. Although those output signals can vary dramatically for different cell types, the basic sensory machine appears to have many common elements. For example, the receptor-like-protein-tyrosine-phosphatase alpha (RPTPα) is needed for rigidity sensing in both fibroblasts and neurons but in hippocampal neurons the loss of rigidity sensing results in long straight neurites whereas in fibroblasts it results in transformed growth^32^. Since DAPK1 is part of rigidity sensing modules in fibroblasts, those cells can activate apoptosis on soft surfaces whereas DAPK1 is not linked to matrix rigidity when the sensory modules are missing cytoskeletal components and do not assemble as in cancer cells. Signaling from Src and RTKs is part of the rigidity-sensing module and that is needed to inactivate DAPK1 as well as activate growth because of force-dependent, tyrosine phosphorylation on rigid substrates. The competing tyrosine phosphatases such as PTPN12 and possibly LAR are designed to keep cells from growing inappropriately when the matrix is soft but are not operative in the absence of the rigidity sensing modules. This raises many questions about how DAPK1 is inactivated when the rigidity sensors are missing and how are the phosphatases recruited to the adhesions on soft surfaces.

## Materials and Methods

### Cell culture

Mouse embryonic fibroblast cells (MEFs) were generated by J. Sap’s laboratory^33^. MDA-MB-231 and MCF10A cells were obtained from J. Groves (University of California, USA and Mechanobiology Institute, National University of Singapore, Singapore). The talin1−/− fibroblast cell line dj26.28 was generated and maintained as described previously^34^. The MEFs and MDA-MB-231 cells were cultured in Dulbecco’s Modified Eagle Medium (DMEM) (Thermo Fisher Scientific) supplemented with 10% fetal bovine serum (FBS) (Atlanta Biologicals) and 100 IU ml^−1^ Penicillin-Streptomycin (Sigma) at 37°C and 5% CO_2_. MCF10A cells were cultured at 37°C in a 5% CO_2_ incubator in DMEM (Thermo Fisher Scientific) supplemented with 20 ng ml^−1^ EGF (Thermo Fisher Scientific), 10 ng ml^−1^ bovine insulin (Sigma), 500 ng ml^−1^ hydrocortisone, 5% horse serum albumin (Thermo Fisher Scientific), and 100 IU ml^−1^ Penicillin-Streptomycin (Sigma). Imaging experiments were conducted using starvation medium without phenol red or in Ringer’s buffer (150 mM NaCl, 5 mM KCl, 1 mM CaCl_2_, 1 mM MgCl_2_, 20 mM Hepes, and 2 g/L glucose, pH 7.4). Pharmacological inhibitors were as follows: DAPK1 inhibitor (100 nM, EMD Millipore), EGFR inhibitor Gefitinib (10 nM, Santa Cruz), PP2 (200 nM, abcam), calpain Inhibitor ALLN (100 μM, Santa Cruz), PTPN12 inhibitor (1.5 μM, EMD Millipore). Cells were trypsinized using TrypLE (Thermo Fisher Scientific) on the following day and suspended in Ringer’s buffer at 37°C for 30 minutes prior to plating on human plasma fibronectin (10 μg/ml, Roche) coated glass-bottom dishes (MatTek) or pillars. The 0.2 kPa PDMS gels used for testing cells’ growth on soft matrices were purchased from Soft Substrates^TM^.

### Transfection and plasmids

Transfections were carried out 1 day before measurements using Lipofectamine Plus Reagent (Invitrogen) according to the manufacturer’s instructions. Expression vectors encoding the following fluorescent fusion proteins were used: EGFP-DAPK1 WT, EGFP-DAPK1 K42A, mCherry-DAPK1 WT, mCherry-DAPK1 K42A, YFP-Tpm2.1 WT, YFP-Tpm2.1 S283E (phospho-mimic mutant), YFP-Tpm2.1 S283A (non-phosphorylated mutant), mCherry-paxillin, GFP-talin1 WT, mRuby-talin1 head, mCherry-talin1 rod, GFP-non-cleavable talin1, EGFP-DAPK1-DYD (phospho-mimic mutant), EGFP-DAPK1-DYF (non-phosphorylated mutant), EGFP-DAPK1-DYF-K42A (non-phosphorylated inactive mutant), mCherry-PTPN12. Cells were seeded into a 6-well dish on day 0 and transfected with control shRNA (Sigma) or DAPK1 shRNA (Sigma) using Lipofectamine Plus Reagent (Invitrogen) on day 1, followed by selection in 2 μg/ml puromycin for 72 hours.

### Pillar fabrication

Fabrication of pillar arrays with diameters of 500 nm was performed as previously reported^24^. Polydimethylsiloxane (PDMS, Sylgard 184, Dow Corning) was mixed thoroughly with its curing agent (10:1) for at least 5 minutes, centrifuged (2000 RPM, 3 minutes, room temperature) and degassed in a vacuum for about 10 minutes. A drop of PDMS was then placed on the top of the Poly (methyl methacrylate) (PMMA) mold and degassed again in a vacuum for a few minutes. At the same time, a glass-bottom petri dish (No.0 Coverslip, MatTek) was treated with O2 plasma for 1 minute, in order to increase adhesion of the PDMS to the glass after curing. The mold with PDMS was then inverted and placed onto the bottom of the petri dish, and a 1 gram weight was placed on the top of mold to make the PDMS thinner. The PDMS was then cured at 70°C for 12-14 hours, in order to achieve a Young’s modulus of 2 ± 0.1 MPa. The mold was then peeled off the PDMS while immersed in ethanol. Individual pillars were 500 nm in diameter with a center to center distance of 1μm.

For the dual stiffness pillar fabrication, Coumarin 343 (Sigma-Aldrich), which will be bleached under UV treatment, was premixed in PDMS for the marking of areas with different rigidity. The dyed PDMS was applied to the master molds, which were spun at 6600 RPM for 3 seconds to ensure even coating. After the PDMS was cured at 70°C for 12-14 hours, it was carefully peeled off the molds and placed (pillar-side up) onto the O_2_ plasma-treated glass-bottom petri dish. Prior to UV/Ozone treatment, nickel TEM grids (SPI supplies) were laid on the pillar tops. Pillar substrates covered by grids were then placed in the UV/Ozone chamber (Bioforce Nanosciences UV/Ozone ProCleaner Plus) for 2 hours.

### Traction force measurements

Cells were spreading on pillar arrays coated with fibronectin (10 μg/ml, Roche). Time lapse imaging of pillars was performed with bright-field microscopy using an Orca-flash 2.8 camera (Hamamatsu) attached to an inverted microscope (Olympus IX81) maintained at 37° C running MicroManager software (UCSF). Images were recorded at 1 Hz using a 100x objective (1.4 NA oil immersion, Olympus). Videos were processed with ImageJ (National Institutes of Health) using the Nano Tracking plugin to track the position of pillars. The time-series positions of all pillars in contact with the cell were fed into a MatLab program (MathWorks) to generate displacement maps as explained previously^17,19^.

### Immunostaining and fluorescence microscopy

Cells were fixed for 15 minutes with 4% formaldehyde solution (Sigma, diluted from 37% in PBS). Fixed cells were then permeabilized with 0.2% Triton X-100 in PBS (V/V, Sigma) for 5 minutes, and blocked in 2% bovine serum albumin (BSA) in PBS (W/V, Sigma) at room temperature for one hour. Primary antibody was diluted in 1% BSA and incubated on samples overnight at 4° C. Secondary antibody was diluted in 1% BSA and incubated on samples for 1 hour at room temperature. For immunostaining, the following primary antibodies were used: anti-paxillin (Abcam, dilution 1:250), anti-DAPK1 (BD Biosciences, dilution 1:50), anti-Tpm2.1 (Abcam, dilution 1:1000), anti-pSer308 DAPK1 (LSBio, dilution 1:100), anti-Annexin V (Abcam, dilution 1:100). Secondary antibodies were Alexa Fluor 488 goat anti-mouse IgG (H+L) and Alexa Fluor 555 goat anti-rabbit IgG (H+L) (Thermo Fisher Scientific).

Total internal reflection fluorescence (TIRF) images were taken using an Olympus IX81 fluorescence microscope with a 60x, 1.45 NA oil-immersion objective and a Cool Snap FX cooled CCD camera (Photometrics) controlled by MicroManager software (UCSF). Confocal microscopy was performed on a Zeiss LSM 700 laser-scanning confocal microscope, using a 63x/1.4 NA oil-immersion objective. Quantification of images was performed with ImageJ (National Institutes of Health).

For the fluorescence recovery after photobleaching (FRAP) experiment, MEFs transfected with EGFP-DAPK1 K42A or EGFP-DAPK1 WT were allowed to spread on fibronectin-coated pillars for 20 minutes and then the focal adhesion areas were bleached using fast bleach on Zeiss LSM 700. Fluorescence intensity recovery was simultaneously recorded at 1 frame per 2 seconds. The images were then corrected for background using ImageJ (National Institutes of Health). Single exponentials were then fitted to the recovery profiles in Excel and SigmaPlot 12.0 software (Systat Software Inc.) to calculate half-times (t_1/2_).

### Statistical analysis

Statistical analysis and graph plotting were performed using SigmaPlot 12.0 software (Systat Software Inc.) and Matlab (Math Works). Two-tailed Student’s t test was performed when two cases were compared. One-way-ANOVA was performed for multi-group comparison. P values <0.05 were considered statistically significant.

## Acknowledgements

We thank all the members of Sheetz lab for their kind help. This work was funded by National Institutes of Health (NIH) grant “Analysis of 120 nm local contractions linked to rigidity sensing” (1 R01 GM100282-01), National Institutes of Health (NIH) grant “Tropomyosin and tyrosine kinases in mechanics of cancer” (5 R01 GM113022-02). H.W. was supported by a Marie Curie International Outgoing Fellowship within the Seventh European Commission Framework Programme (PIOF-GA-2012-332045), and is an incumbent of the David and Inez Myers Career Advancement Chair in Life Sciences. M.P.S is supported by NUS grants and MBI, National University of Singapore.

## Author Contributions

R.Q. H.W. M.S. and M.P.S conceived the study and designed the experiments; R.Q. and H.W. performed the experiments; R.Q. and H.W. analyzed the data; R.Q. H.W. and M.P.S wrote and prepared the manuscript.

